# OrchID: a Generalized Framework for Taxonomic Classification of Images Using Evolved Artificial Neural Networks

**DOI:** 10.1101/070904

**Authors:** Serrano Pereira, Barbara Gravendeel, Patrick Wijntjes, Rutger A. Vos

## Abstract

Taxonomic expertise for the identification of species is rare and costly. On-going advances in computer vision and machine learning have led to the development of numerous semi- and fully automated species identification systems. However, these systems are rarely agnostic to specific morphology, rarely can perform taxonomic “approximation” (by which we mean partial identification at least to higher taxonomic level if not to species), and frequently rely on costly scientific imaging technologies. We present a generic, hierarchical identification system for automated taxonomic approximation of organisms from images. We assessed the effectiveness of this system using photographs of slipper orchids (Cypripedioideae), for which we implemented image pre-processing, segmentation, and colour and shape feature extraction algorithms to obtain digital phenotypes for 116 species. The identification system trained on these digital phenotypes uses a nested hierarchy of artificial neural networks for pattern recognition and automated classification that mirrors the Linnean taxonomy, such that user-submitted photos can be assigned a genus, section, and species classification by traversing this hierarchy. Performance of the identification system varied depending on photo quality, number of species included for training, and desired taxonomic level for identification. High quality photos were scarce for some taxa and were under-represented in the training set, resulting in imbalanced network training. The image features used for training were sufficient to reliably identify photos to the correct genus but less so to the correct section and species. The outcomes of this project include a library of feature extraction algorithms called *ImgPheno,* a collection of scripts for neural network training called *NBClassify,* a library for evolutionary optimization of artificial neural network construction called *AI::FANN::Evolving* and a planned web application called *OrchID* for identification of user-submitted images. All project outcomes are open source and freely available.

## INTRODUCTION

Correct taxonomic identification of organisms is of great importance. However, taxonomic expertise is rare and costly (the “taxonomic impediment”[1]), so alternative, more scalable methods that are cheap, that require little expert knowledge, and that are accurate enough to derive taxonomic approximations – i.e. partial identification at least up to higher taxonomic levels such as genera, though under ideal circumstances to species level – would be of great value. Taxonomic identification and approximation using DNA barcodes (for example, to detect biological materials from endangered species [2]) may be dropping in relative cost and may require little taxonomic expertise, but the need for wet lab procedures and DNA sequencing is still prohibitive in many contexts. Low-cost taxonomic identification systems based on geometric morphometrics (for example of bee wing morphology [3]) require human assistance to identify homologous landmarks, which at least requires expertise in morphology, in addition to the time-consuming handwork of applying landmarks, which impedes scalability.

Considering the case of slipper orchids (Cypripedioideae), a popular horticultural group of orchids of which many species are highly endangered in the wild and in which illegal trade is soaring [5,12], more tools are needed to increase taxonomic identification to prevent species from going extinct. On-going advances in computer vision and machine learning have led to the development of numerous semi-and fully automated species identification systems. Such systems have been successful in the identification of plant species [4–6], phytoplankton [7], diatom frustules [8], and insects [9,10], among others. However, these systems frequently rely on costly scientific imaging and analysis (e.g. microscopy-based histology [6,8] or flow cytometry [7]) and rarely perform taxonomic approximation, rather taking a binary “all or nothing” approach. An informative exception to this is the work of Boddy et al. [7], who devised a hierarchy of classifiers whose structure mirrors that of the Linnean taxonomy such that queries are passed on from classifiers at higher taxonomic levels to the appropriate classifier at lower levels. This approach allows for taxonomic approximation, and it is enabled by the application of Artificial Neural Networks (ANNs). ANNs have become an increasingly popular choice in automated image classification systems [12]. They provide a powerful tool for pattern recognition and machine learning. However, due to the frequently very large number of configuration parameters (for network topology, activation functions, training algorithms, and so on), ANN construction and training is difficult to generalize.

Table 1 lists some of the recent studies that present systems for classifying organisms from images. Although some studies are based on (partially) manually applied landmarks [3,6], the majority of these systems use various ways to automatically capture and summarize putatively informative image features, e.g. by computing colour intensities and/or shape descriptors. The image data are obtained from a variety of technologies, about half of which are rely on standard, consumer-quality cameras, with the others using various types of microscopy or other ways of detecting microscopic features (i.e. by flow cytometry, [7]). Although the number of different statistical methods used to arrive at classifications is striking, the most commonly used approach is based on ANNs. Interestingly, few systems attempt to arrive at taxonomic approximation, and those that do in our survey [7,10], rely on specialized data capture technologies.

**Table 1.** Prior art in image recognition and classification of organisms. KDA=Kernel Discriminant Analysis; SVM=Support Vector Machine; DFA=Discriminant Factor Analysis; ANN=Artificial Neural Network; EFA=elliptic Fourier analysis; LDA=Linear Discriminant Analysis. “Summary” image features are those that are computed by summarizing one or more image properties, such as colour intensities or the properties of an object outline.

| **Project name** | **Image type** | **Classifier type** | **Feature type** | **Taxonomic approximation** | **Literature reference** |
| --- | --- | --- | --- | --- | --- |
| ABIS | Microscopy | KDA | Landmarks | No | [3] |
| VISIONTRAIN | Standard | SVM | Summary | No | [4] |
| Sanz et al. | Standard | DFA | Summary | No | [5] |
| Arinkin et al. | Microscopy | ANN | Landmarks | No | [6] |
| Boddy et al. | Cytometry | ANN | Summary | Yes | [7] |
| SHERPA | Microscopy | EFA | Summary | No | [8] |
| Kang et al. | Standard | ANN | Summary | No | [9] |
| DAISY | Microscopy | Kendall-τ | Summary | Yes | [10] |
| Corney et al. | Standard | LDA | Summary | No | [11] |

Given the current state of the art we propose that generalizable systems for taxonomic approximation, agnostic to specific morphology, and applicable to low-cost imaging equipment such as phone cameras, should be coming within reach of non-expert users. We therefore present a novel framework to fill this void. The framework comprises reference image management; image pre-processing, segmentation, and naïve, unassisted feature extraction; and training of hierarchically nested ANNs. We address the challenges of ANN parameterization using an evolutionary algorithm, and bundle all functionality to classify user-submitted photos as a handheld-compatible web application. To demonstrate the feasibility of the approach we apply it to taxonomic approximation within the slipper orchids (Cypripedioideae), a charismatic group of wild plant species whose exuberant flower shapes and colours make them very popular ornamentals among gardeners, orchid enthusiasts and wildlife photographers. Many species are highly endangered in the wild due to on-going illegal trade worldwide such that correct taxonomic identification is also necessary to improve identification of confiscated material by a wide variety of non-specialist law enforcers such as customs officers and park wardens.

## MATERIALS AND METHODS

### Reference image management

ANNs must be trained with a sufficient number of exemplars to inform classification of new, out-of-sample queries. We therefore compiled a collection of reference photos by searching the Internet for images, mostly through Google Image searches. We obtained images from several multimedia resources such as Flickr (https://www.flickr.com) and Wikimedia Commons (http://commons.wikimedia.org), but also from botanical gardens, nurseries, orchid foundations and societies and wildlife photographers (see Acknowledgements). Given the progressively growing size of the reference image set and the amount of metadata on taxonomic classification and image provenance that needed to be tracked, we required a simple but consistent strategy for collaboratively managing all this. An ideal solution would be a central server where image sets can be assembled along with their metadata (i.e. taxonomic information required for training the neural networks, and provenance data on the photo per se). Such a database needs to be accessible via an application-programming interface (API) that allows for the implementation of custom image harvesters for easy retrieval of images and their metadata.

We used the online image hosting service Flickr since it meets these requirements. We uploaded photos to an account on Flickr, where we annotated them with standardized tags of the form *genus:<name>*, *section:<name>*, and *species:<name>*. Here, *section* is understood to refer to its usage in botanical taxonomy, i.e. as a level in the Linnean hierarchy between genus and species. We verified the correct genus, section, and species for each training image with specialist taxonomic knowledge (BG), literature study [13–15] and phylogenetic reconstructions based on molecular analyses [16,17]. Only wild species were used so all horticultural hybrids were left out. We implemented a custom Python script utilising the Flickr API to mirror these reference image sets and associated metadata in a local file store prior to analysis. We stored images in an automatically generated directory hierarchy such that the nested directory names correspond to the taxonomy. In addition, we created a local relational database (Supplementary Figure 1) to store and query metadata for each reference image set, as this is needed in downstream image processing.

### Image pre-processing and feature extraction

As part of the image pre-processing and feature extraction workflow, each image is first scaled down if the image exceeds a predefined maximum dimension. This is followed by foreground segmentation to get the region of interest (ROI) that matches, in this case, the flower. Foreground segmentation is done with an iterative approach using the GrabCut segmentation algorithm [18]. The ROI for the first iteration is set to the entire image and is shrunk a fixed number of pixels to get a margin around the ROI containing obvious background pixels (Fig. 1). Images must therefore be reasonably standardized such that the flower is within this initial ROI. There must also be good contrast between the flower and the background, as to keep segmentation errors to a minimum. Subsequent iterations further classify pixels inside the ROI as foreground or background until the maximum number of iterations is reached. Contours are then obtained from the resulting binary mask, which is a copy of the image where foreground pixels are made white and background pixels black. Sometimes multiple contours are found (e.g. on images with multiple flowers) in which case only the largest contour is used to select the foreground pixels for downstream feature extraction.

**Figure 1.**
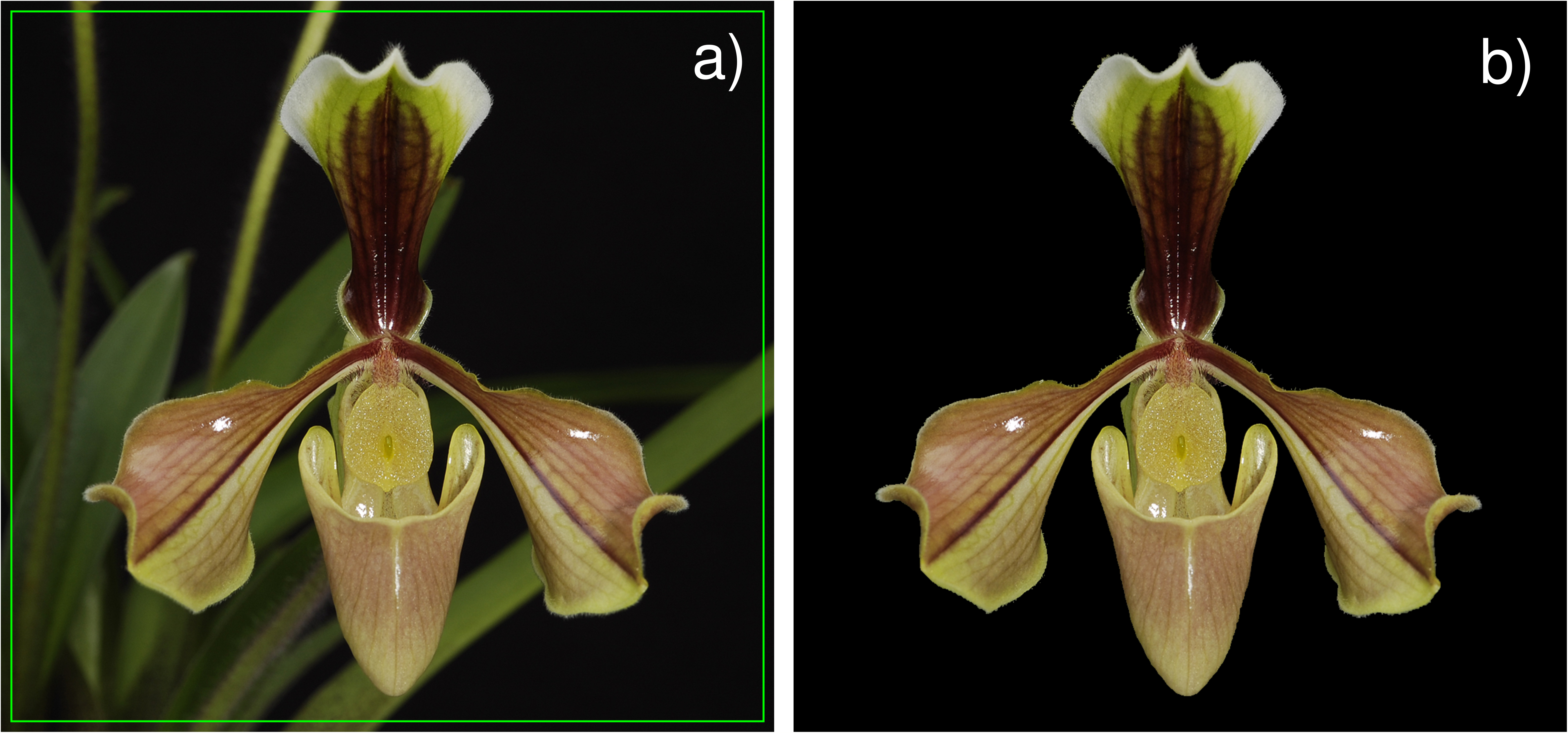
Foreground segmentation of a photo of *Paphiopedilum villosum* using the GrabCut algorithm. **(a)** Shows how the initial region of interest (ROI) is set; **(b)** shows the result of the segmentation using the GrabCut algorithm. Reprinted from http://www.pbase.com/rogiervanvugt/image/109985904 under a CC BY license, with permission from Rogier van Vugt, original copyright 2009.

Since two-dimensional pixel data are usually not appropriate for pattern recognition algorithms (which normally expect as input a fixed-length sequence of numbers) images need to be summarised to one or more numerical values. These values, often called features, describe some aspect of the image contents. The algorithms or applications that compute these features are often called feature descriptors or feature extractors. Many feature extraction algorithms are generic and are not limited to specific image types. Feature extractors can be applied to entire images and sometimes to regions of interest (ROI) in images. Many feature extraction algorithms exist and are described in the literature, and some have been implemented in standalone software or libraries. Image recognition software packages that use pattern recognition and machine learning usually implement one or more feature extraction algorithms to summarize features and use those to train classifiers. With the many different feature extraction algorithms already available and new ones emerging periodically, finding the right algorithm for a specific imageprocessing problem can be a daunting task. Open Source Computer Vision (OpenCV) and Open Intelligent Multimedia Analysis for Java (OpenIMAJ) are comprehensive computer vision libraries that come with computer vision and machine learning algorithms (amongst others), but support for feature descriptors is currently limited. To remedy this, we developed a general purpose, open-source software library of image feature description algorithms for the Python programming language – similar to what JFeatureLib does for Java – called *ImgPheno* (https://github.com/naturalis/imgpheno), which we used as one of the modules in our framework. *ImgPheno* makes extensive use of OpenCV [19] for computer vision, NumPy [20] for array manipulation, and of scikit-learn [21] for data transformation and cross-validation.

We automated feature extraction and subsequent neural network training with a custom Python script that is part of the *NBClassify* package (https://github.com/naturalis/nbclassify). Configurations are kept in a separate YAML file so that neural network training can easily be reproduced. Different aspects of the feature extraction process can be configured with this configuration file: input/output format, data scaling, colour correction, foreground segmentation, feature descriptors to use, and a classification hierarchy definition for hierarchical training.

In the present study we used only one feature extractor to generate data for subsequent training of the artificial neural networks. This feature extractor computes summarised histograms from the pixel colour intensities for the BGR colour space. This is done by calculating the upright bounding square for the main contour, dividing the square into *N* equal horizontal and vertical sections (bins), and calculating the mean blue, green, and red colour intensity for each bin. This results in two histograms, one for the horizontal and one for the vertical bins (Fig. 2). Plotting the histograms shows that this feature captures aspects of flower shape as well (Fig. 3).

**Figure 2.**
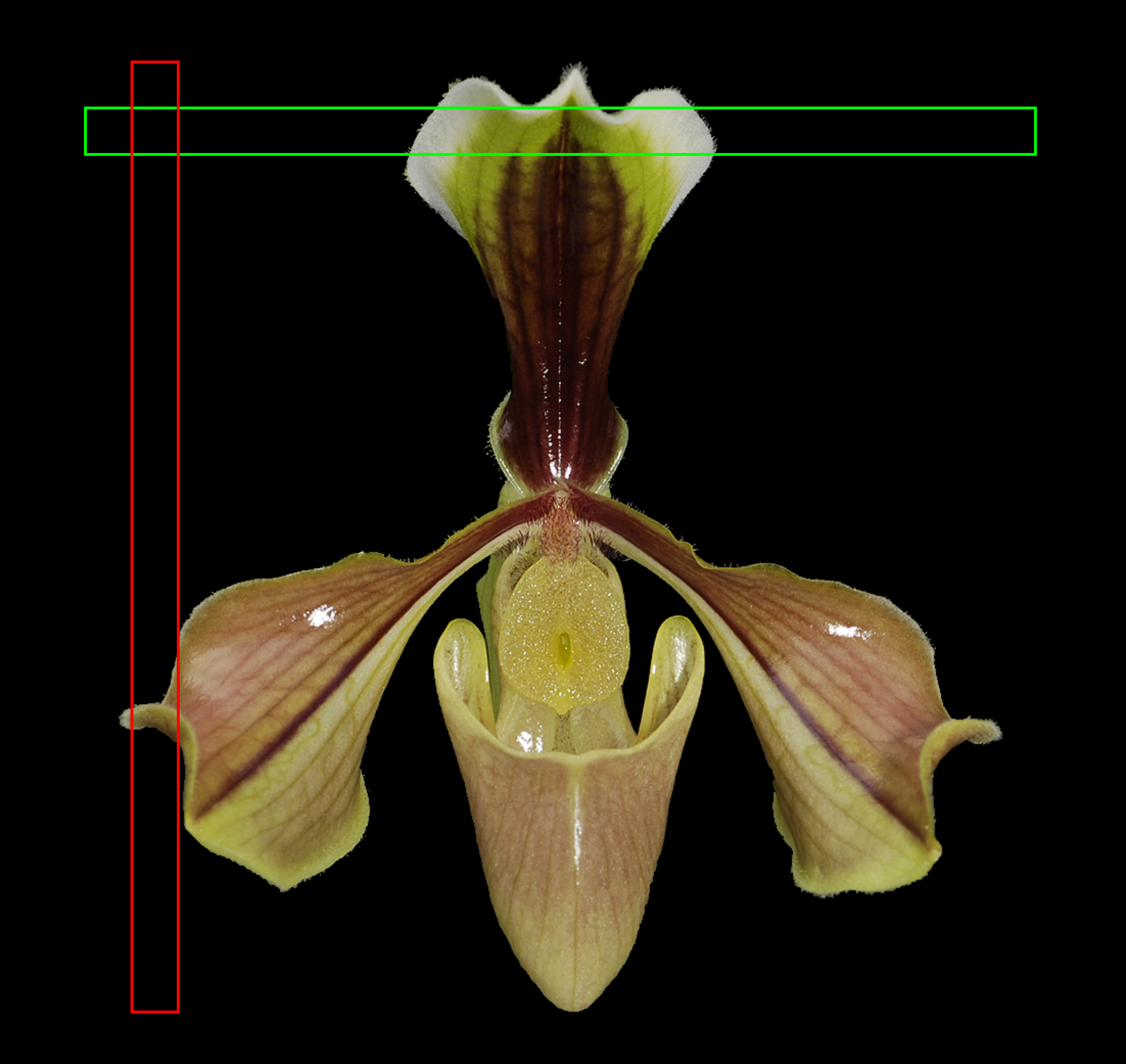
The second horizontal and vertical bin (*N* = 20) highlighted in a foreground-segmented image of *Paphiopedilum villosum.*

**Figure 3.**
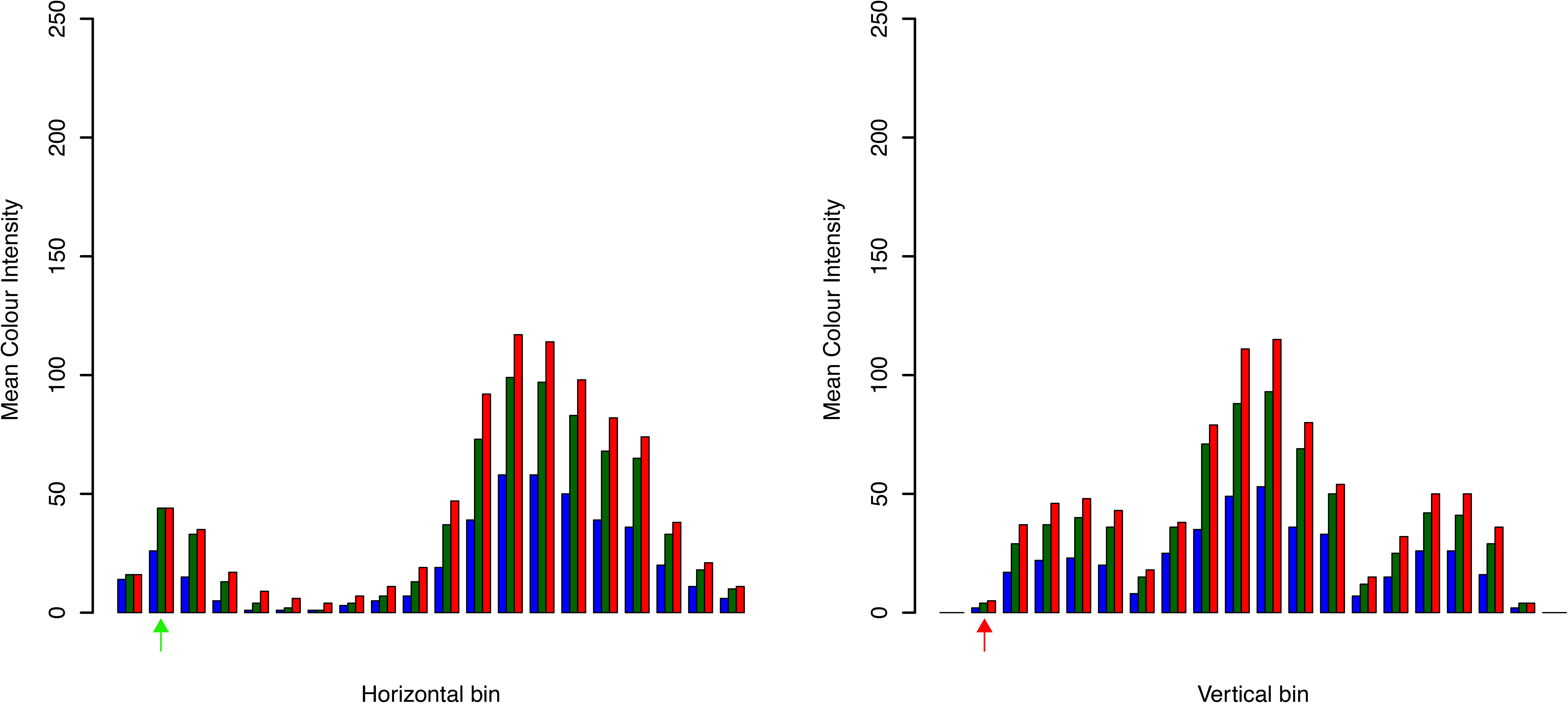
Plots of the mean BGR colour intensities for an image of *Paphiopedilum villosum*. The plots display the mean intensities for the horizontal and vertical bins respectively. Arrows indicate the locations of the bins shown in Figure 2.

We performed principal component analysis (PCA) on the feature data to assess their suitability for differentiating photos by genus, section, and species. We conducted PCA with orthogonal rotation (varimax) on several training data sets containing -1 to 1 scaled mean colour intensities for the BGR colour space with 20 horizontal and vertical bins, which translates to 120 features. We used the Kaiser-Meyer-Olkin measure to verify the sampling adequacy for the analysis. We used Bartlett’s test of sphericity to get indications whether correlations between items were sufficiently large for PCA. We used the inflexion in scree plots to set the number of components to be extracted. We performed PCA on training data for genus classification within the Cypripedioideae, section classification within the genus *Paphiopedilum*, and species classification within section *Parvisepalum* of genus *Paphiopedilum.* We did not perform PCA for the remaining data sets because the number of features often exceeds the sample size.

### ANN training and cross-validation

Because the 140+ species of slipper orchids pose challenges for training a single ANN (essentially, too many classes among which to discriminate), and because a single ANN cannot provide approximation at least to higher taxonomic levels, we used a hierarchical approach. We implemented artificial neural network (ANN) training using the Fast Artificial Neural Network Library (FANN) [22]. We performed hierarchical multi-species classification with a system of feed forward artificial neural networks that were trained using the iRPROP backpropagation training algorithm. Hierarchical training results in multiple neural networks: one ANN for genus classification, one ANN per genus for section classification, and one ANN per section for species classification. For the individual neural networks this means they need to differentiate between fewer classes and they can be trained using rank or taxon specific features. No ANNs need to be trained on nodes where no branching occurs in the hierarchy (e.g. monotypic genera).

In training ANNs there is the risk of overfitting: if an ANN is provided a limited set of training data it may end up constructing responses to the specific idiosyncrasies of these data rather than on more appropriate, general properties that would be recognized had the training data set been larger, or more representative. When testing such overfitted ANNs, for example by presenting them with data from the training set, the problem will manifest in the form of spuriously inflated performance indicators. To mitigate against this we never tested ANNs by presenting them with the same data that were used for training. Throughout this manuscript, we indicate the distinction between training data and test data by referring to the latter as “out-of-sample”, i.e. outside of the training data sample.

Hierarchical classification means that an incorrect classification at a higher rank in the hierarchy propagates to incorrect subsequent classifications at lower ranks, but it also allows for partial classifications when classification fails for the lower ranks, i.e. taxonomic approximation. The problem of propagating erroneous classifications can be mitigated against, at least to some extent, by selecting a stringent threshold value for the ANN’s uncertainty in its output (in FANN terminology, the mean squared error) such that further propagation ceases if the classification is too uncertain.

The classification hierarchy is defined in the configuration file as a list of levels that tells the training script the path to follow during training and classification. In this study the levels were named after the corresponding taxonomic ranks: genus, section and species. Different features and training parameters can be set at each level in the classification hierarchy, which the script then uses to generate training data for the required ANNs. We scaled all training data to a -1 to 1 range as per the span of the used activation function (symmetric sigmoid). We used bit vector-like output sequences to encode the known taxon for each image in the training data. Each ANN has its own set of sequences generated from the sorted lists of taxa the ANN is trained on. Each sequence consists of *N* bits, where *N* is the number of classes the ANN is trained on. All bits in the sequence are set to an off value (-1), except for one bit (1), which corresponds to the class in the sorted list of classes. Once the training data are generated they are used to train the individual neural networks.

FANN supports many training parameters (e.g. training algorithms, activation functions, neural network topology, etc.), making it hard to predict which parameters and which combination of algorithms work best for a given classification case under the generally naïve approach we take (i.e. not tuned to a specific taxonomic group or feature set). We therefore implemented a framework for evolving optimal neural networks (*AI::FANN::Evolving*, https://github.com/naturalis/ai-fann-evolving) to overcome this issue. The evolutionary algorithm models a constant-sized population of diploid, one-chromosome organisms whose crossover and mutation rates can be configured. The algorithm then proceeds for a pre-defined number of generations. At each generation, each individual crosses over and mutates its FANN configuration parameters, trains on a data set, and then tests its classifiers (two: one on each of the homologous chromosomes) on an out-of-sample data set. The individuals are then mated with each other in proportion to the success of their classifiers until the population size for the next generation is reached. Once the pre-configured number of generations is reached, the “fittest” overall ANN is selected.

We performed stratified k-folds cross-validation to estimate the accuracy of hierarchical classification. We used genus-section-species combinations as the classes for the cross-validations. Because the amount of images for some taxa was very limited, we excluded taxa for which the number of images was less than the number of folds *(k).* We performed cross-validation with both standard training and training with optimisation of ANN parameters by the evolutionary algorithm.

## RESULTS

### Reference image collection

We collected a total of 1136 photos for 116 species of slipper orchids from various sources (Supplementary Table 1). The collection represents all five genera from subfamily Cypripedioideae: *Cypripedium, Mexipedium, Paphiopedilum, Phragmipedium,* and *Selenipedium.* With the number of photos for the five genera ranging from just four for the genus *Selenipedium* to 888 for *Paphiopedilum,* the collection is highly unbalanced. There is also a high variation in the number of images within genera. Unbalanced data results in a bias towards the more image-rich taxa during neural network training, meaning the under-represented taxa are less likely to be classified correctly.

Because images of some taxa were hard to find, we had to make some compromises with respect to the quality of the images. Many of the gathered images are not of the desired standardised format, resulting in variations in background, lighting, dimension, and flower position or rotation, which makes pattern recognition more challenging. Since foreground segmentation is part of the automated workflow, this sometimes results in incorrectly segmented images.

We have stored the complete collection of reference images on Flickr (https://www.flickr.com/photos/113733456@N06/). However, as these images were supplied by a variety of different contributors (listed in the Acknowledgements section), who have varying requirements for attribution and reproduction, we have had to store our reference images in a password-protected collection. Access to this collection will be granted to all researchers interested in using these images for research purposes, provided the images are not reproduced without permission from the original copyright holders. Once access is granted, reference images can be downloaded in batch and placed in a directory hierarchy using the *nbc-harvest-images* script of the *NBClassify* package. In addition, the *nbc-trainer* script can be used to create a meta-data database from an existing image directory, which would enable subsequent training on images of that directory.

### Image pre-processing and feature extraction

The *ImgPheno* library is currently at an early development stage and only a few feature extraction algorithms have been implemented so far. The library also includes implementations of colour enhancement algorithms, helper functions for feature drawing, and algorithms that implement image features to compute shape properties (e.g. computing the centre point from moments). The intent of the *ImgPheno* library is to provide a generic toolkit for image processing and feature extraction that can be applied to problems beyond the scope of the present study. As such, *ImgPheno* contains functions beyond those strictly required for our current work. For example, although we have implemented a number of functions operating on object contours, we found that they do not aid in discriminating among classes of orchid flowers (results not shown). An overview of the descriptors and enhancement functions can be found in Table 2.

**Table 2.** *ImgPheno* features and methods overview.

| Method | Description |
| --- | --- |
| Colour Histogram | Returns the histogram for an image. |
| Binned Colour Histogram | Returns histograms for the mean BGR intensities from the horizontally and vertically binned image. |
| Contour Eccentricity | Computes a scalar that specifies the eccentricity of the ellipse that fits the contour. The eccentricity is the ratio of the distance between the centre and either focus of the ellipse and its major axis length. |
| Contour Equivalent Diameter | Computes a scalar that specifies the diameter of a circle with the same area as the contour. |
| Contour Solidity | Computes a scalar specifying the proportion of the pixels in the convex hull that are also in the region. |
| Contour Outline | Computes a vector that summarizes the outlines of a contour at a specified resolution. |
| Contour Centre Based Branch Lengths | Computes a vector that summarizes the outline of a contour using branch lengths from the centre of mass. |
| Linear Colour Enhancement | Provides a hue preserving linear transformation with maximum possible contrast [24]. |
| Non-linear Colour Enhancement | Provides non-linear hue preserving transformation without gamut problem, provided that linear transformation is initially applied on each of the pixels [24]. |
| S-type enhancement function | This implements the S-type enhancement function for contrast enhancement of grey scale images [24]. Can be used as the enhancement function for non-linear colour enhancement. |

Figure 4 shows the scatter plots for the principal component analyses (PCAs) that we performed on a subset of the feature data. Poor classification accuracies shown in Table 3 (discussed in more detail below) may be attributed to the poor discrimination ability of the colour-based feature to the different taxa and to improper foreground segmentation. Out of the four plots, only the plot for species classification within section *Parvisepalum* of genus *Paphiopedilum* showed some clear clusters. The accuracy for species classification within this section, given correct classification of genus and section, was 85% and 88% for 4 and 10 folds, respectively. This compares fairly well with the accuracy for genus classification (75% and 81% for 4 and 10 folds, respectively), which is surprising when comparing the scatter plots for genus classification and *Parvisepalum* species classification.

**Figure 4.**
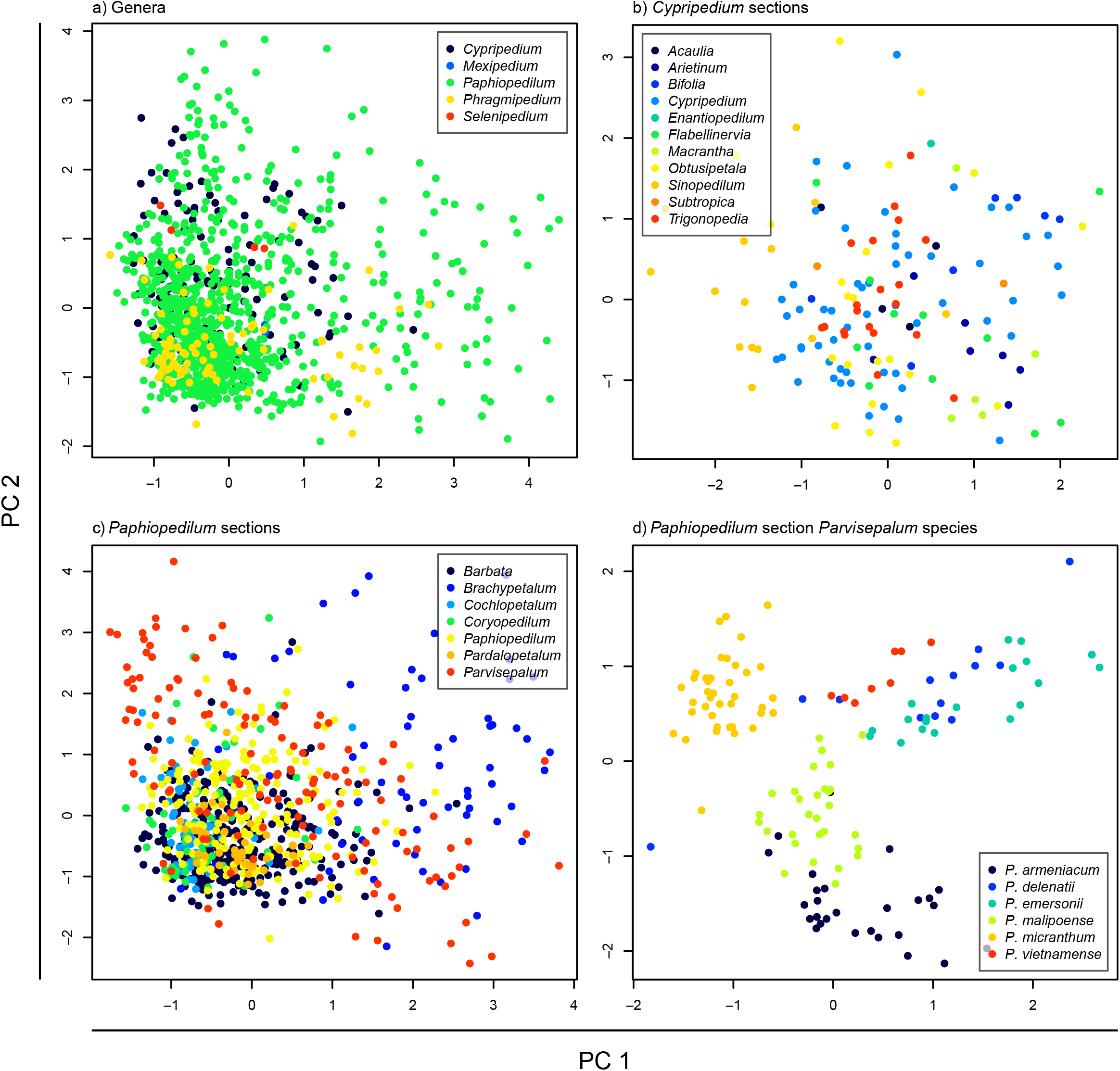
Scatter plots for the principal components analysis results. PCA was conducted on training data for (**a**) all genera; (**b**) all sections within genus *Cypripedium;* (**c**) all sections within genus *Paphiopedilum*; and (**d**) all species within *Paphiopedilum* section *Parvisepalum*. For all plots the first principal component (PC1) was plotted against the second (PC2).

**Table 3.** Results for stratified k-fold cross-validations. Cross-validation was performed on three taxonomic ranks: genus, section, and species. The results for genus/section and genus/section/species nest the results from their respective ranks.

| **Classification** | **Accuracy (k=4)** | **Accuracy (k=10)** |
| --- | --- | --- |
| Genus | 75% | 81% |
| Section | 52% | 60% |
| Species | 48% | 56% |
| Genus/section | 41% | 49% |
| Genus/section/species | 20% | 27% |

### ANN performance cross-validation

We performed stratified k-folds cross-validation to estimate the accuracy of hierarchical classification. The accuracy for genus classification was 75%, but accuracy drops as section (52%) and species (48%) are also included in the classification (Table 3). With 10 folds and including only those species for which at least 10 photos were collected, we achieved slightly better accuracies, which can probably be attributed to the reduced number of taxa and more equal image representation between taxa.

All results of the cross-validation are available in the data repository of the project (doi:10.5281/zenodo.31904) which includes the following: the phenotypes extracted from the input images (i.e. the actual input data); a placeholder directory that can be readily populated with the reference images using a script that harvests these from the Flickr library provided valid login credentials (we resort to this approach because some images shared with us cannot be posted publicly); all ANNs that were trained during cross-validation; the training and testing data partitions; and the classification results.

### Bundling of interacting modules in the framework

The interaction of the different modules in our framework is visualized in Figure 5. Briefly, reference image management is an asynchronous, collaborative process that uses the Flickr API (Fig. 5a); image pre-processing and feature extraction, both for reference images and user-submitted query images is based on the *ImgPheno* library, while ANN training and evolutionary optimization of ANNs are handled, respectively, by *NBClassify* and *AI::FANN::Evolving* (Fig. 5b); users can submit images for out-of-sample classification through a proof-of-concept web application called *OrchID* (under development in Python using the Django framework, source code included in the *NBClassify* repository; Fig. 5c).

**Figure 5.**
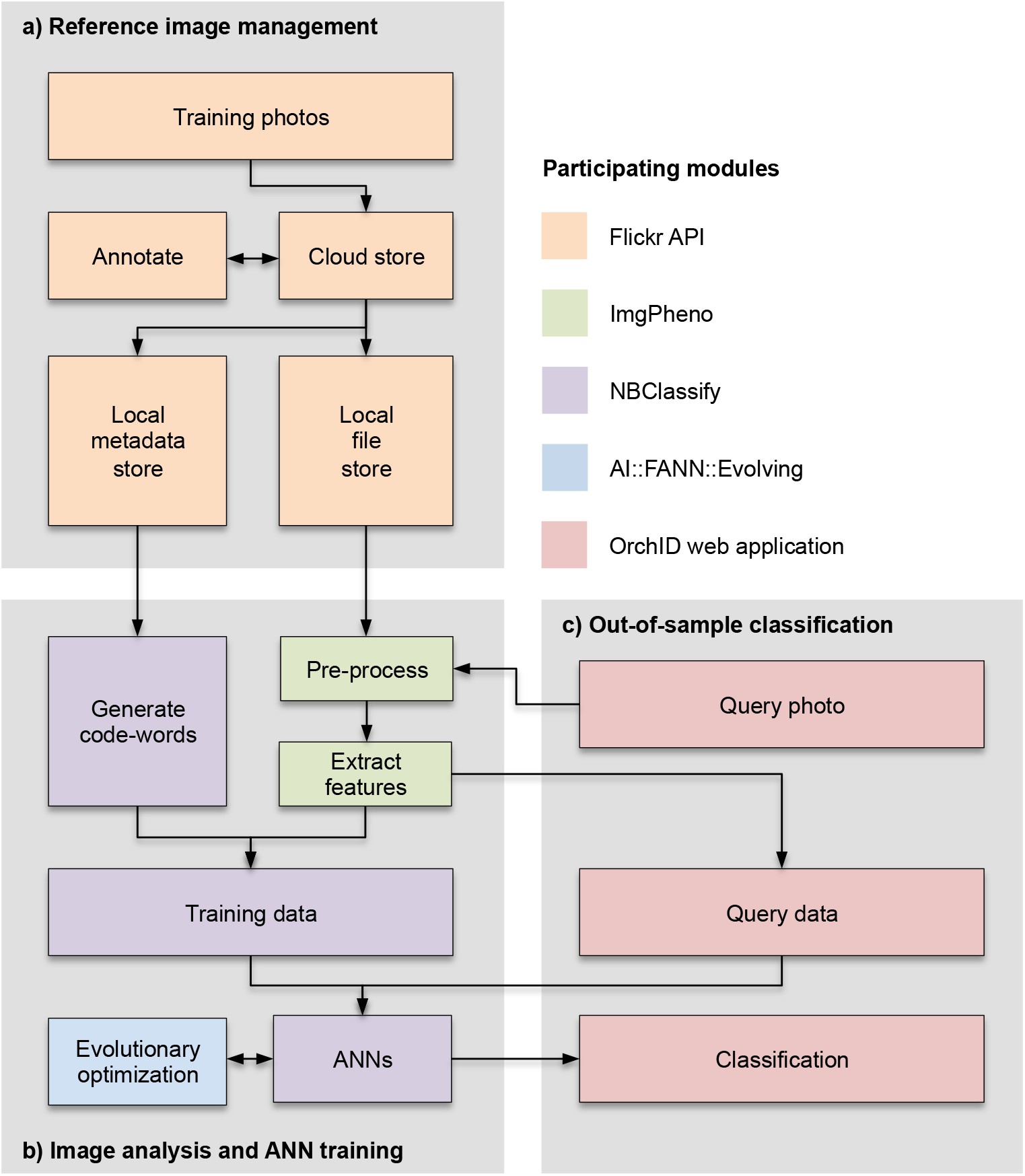
Interactions between modules in the framework. (**a**) Management, annotation and mirroring of reference images using the Flickr API. (**b**) Image pre-processing and feature extraction using *ImgPheno*; ANN training using *NBClassify*; ANN evolutionary optimization using *AI::FANN::Evolving.* (**c**) Out-of-sample classification of user-submitted query images using the *OrchID* web application. The processes in (**a**) and (**b**) are asynchronous, preparatory steps; the processes in (**c**) happen in real time.

## DISCUSSION

Our results indicate that hierarchically nested ANNs trained on naïve image features can perform taxonomic approximation for out-of-sample images of slipper orchids. We obtained our results by extracting simple histograms of colour intensities along the horizontal and vertical axes of (pre-processed) input images, i.e. the image features we extracted are not based on any properties that are specific to orchid flowers. However, we did make certain assumptions about the input images that should also be met by any other group of taxa to which our approach might be applied. These assumptions are that:

i. From the input images, a single region of interest can be selected. We have permitted input images that contained multiple flowers, corresponding with multiple potential regions of interest, but for these we automatically selected the single largest candidate region, i.e. a single flower.
ii. The selected region of interest consists of an object that can be orientated in the same direction consistently across all input images. In the case of the orchid flowers this meant that the input image had to contain a frontal view of a flower, for which we corrected any rotation such that in all pre-processed images the dorsoventral axis is exactly vertical.
iii. The amount of variation of form within each image class is limited. For example, all images in class are from objects in the same developmental stage, and all parts of the objects are more or less in the same place. Indeed, we noticed that the performance of the classifiers was relatively poor in classes that showed a relatively large amount of variation, such as in the case of *Coryopedilum,* where the long, twisted petals can point in different directions.
iv. Using the simple features we extracted, colour must be one of the discriminating features. We expect that other feature descriptors (e.g. variance in intensity among (groups of) pixels so as to capture speckled or striped phenotypes) can be readily and usefully implemented in *ImgPheno* but have not tested these in our current study.

Consequently, the framework should be applicable to (parts of) other symmetrical, regularly shaped, colourful organisms, such as butterflies, beetles, colourful bivalves, and so on. Given the recursive design of our hierarchically nested classifiers, more taxonomic levels can readily be included, such that traversal along the Linnean hierarchy would be extended from higher taxa (e.g. families, or orders).

The framework requires many high-quality reference images, which need to be balanced across classes. For example, the considerably higher accuracy in identifying out-of-sample images as *Paphiopedilum* (Table 4) compared to other genera is probably to a large extent the result of the much larger number of reference images for this genus. This may also explain the higher accuracy in classifying to genus level versus section or species (75% vs. 52% vs. 48%). Likewise, the accuracy of the classification to section level strongly depends on the number of reference images, but is also influenced by variation in shape and colour (Table 5). For example, the lowest number of reference images (13) that still yielded an accuracy of 50% in 10-fold cross-validation was for section *Sinopedilum*, which are very distinct in their brownish colour and compact, rounded petal shape, while 22 reference images still only yielded an accuracy of 13.33% in 10-fold cross-validation for section *Coryopedilum*, which are characterized by their long, twisted petals that extend at variable angles, presumably introducing too much variation in the extracted features. Note that the counts here indicated for reference images (respectively, 13 and 22) do not directly correspond with the raw numbers in Table 5. This is because in our implementation of *k-* fold validation all images are stratified at species level such that all species whose number of available images is < *k* are excluded from validation analyses, also at higher taxonomic levels. The aggregate number of available images in the cross-validation at higher taxonomic levels is consequently lower than the raw counts.

**Table 4.** Number of sections and species per genus, number of photos, and 10-fold cross-validation results at genus level, where applicable. Where too few photos were available for a species (i.e. < k), and where the candidate genus subtends only a single section and/or species, the absence of validation is indicated with ‘not applicable’ (N/A).

| **Genus** | **Sections** | **Species** | **Photos** | **Accuracy** |
| --- | --- | --- | --- | --- |
| *Cypripedium* | 11 | 24 | 149 | 28.45% |
| *Mexipedium* | 1 | 1 | 8 | N/A |
| *Paphiopedilum* | 7 | 71 | 886 | 89.95% |
| *Phragmipedium* | 4 | 19 | 89 | 33.17% |
| *Selenipedium* | 1 | 1 | 4 | N/A |

**Table 5.** Results for k-fold cross-validations at section level. Where too few photos were available for a species (i.e. < k), and where the candidate genus subtends only a single section, and/or species the absence of validation is indicated with ‘not applicable’ (N/A).

| **Section** | **Genus** | **Photos** | **4-fold (nested)** | **10-fold (nested)** |
| --- | --- | --- | --- | --- |
| *Barbata* | *Paphiopedilum* | 326 | 58.65% (54.57%) | 65.62% (61.85%) |
| *Parvisepalum* | *Paphiopedilum* | 135 | 72.66% (56.61%) | 70.46% (60.08%) |
| *Paphiopedilum* | *Paphiopedilum* | 188 | 57.30% (46.91%) | 58.89% (54.57%) |
| *Coryopedilum* | *Paphiopedilum* | 60 | 10.34% (5.21%) | 13.33% (13.33%) |
| *Brachypetalum* | *Paphiopedilum* | 63 | 58.67% (50.85%) | 54.07% (47.21%) |
| *Pardalopetalum* | *Paphiopedilum* | 48 | 30.71% (22.00%) | 20.00% (17.50%) |
| *Cochlopetalum* | *Paphiopedilum* | 66 | 49.58% (43.11%) | 46.83% (43.17%) |
| *Coryopedilum* | *Paphiopedilum* | 60 | 10.34% (5.21%) | N/A |
| *Micropetalum* | *Phragmipedium* | 38 | 50.56% (32.15%) | 100.00% (45.00%) |
| *Phragmipedium* | *Phragmipedium* | 26 | 39.17% (0.00%) | 100.00% (0.00%) |
| *Lorifolia* | *Phragmipedium* | 21 | 46.67% (14.58%) | N/A |
| *Cypripedium* | *Cypripedium* | 57 | 36.74% (13.07%) | 53.33% (6.67%) |
| *Obtusipetala* | *Cypripedium* | 21 | 57.50% (10.00%) | 85.00% (20.00%) |
| *Trigonopodia* | *Cypripedium* | 20 | 37.08% (16.67%) | 40.00% (30.00%) |
| *Sinopedilum* | *Cypripedium* | 14 | 20.83% (14.58%) | 50.00% (15.00%) |
| *Flabellinervia* | *Cypripedium* | 11 | 25.00% (8.33%) | N/A |
| *Bifolia* | *Cypripedium* | 7 | 50.00% (12.50%) | N/A |
| *Arietinum* | *Cypripedium* | 7 | 0.00% (0.00%) | N/A |
| *Acaulia* | *Cypripedium* | 4 | 0.00% (0.00%) | N/A |
| *Macrantha* | *Cypripedium* | 5 | 25.00% (0.00%) | N/A |
| *Selenipedium* | *Selenipedium* | 4 | N/A | N/A |
| *Mexipedium* | *Mexipedium* | 8 | N/A | N/A |

Identification accuracy can probably be improved by means of neural network ensembles where multiple neural networks are used to make classifications [23]. The identification system presented here does use multiple neural networks for the identification of an image, but these cannot be considered neural networks ensembles because only one neural network is used for each level within the hierarchical classification. The output codes used for training the neural networks as implemented in this study is another aspect that could be improved. The implementation presented here uses the simplest possible output codes for training, but the usage of more sophisticated codes could improve neural network training and therefore identification accuracy [24].

Likewise, identification accuracy can probably also be improved by using more, or different, image features. For example, other summary statistics along bins can be computed, such as to capture variance in colour (high variance would then correspond with stripes or dots, while more homogeneous colouration would show lower variance). In addition, analogous to the use of multiple morphological characters in species identification by taxonomic experts, a combination of multiple features may be used for ANN training, potentially with different features being used at different levels of the hierarchical classification system. Developing such specialized features extraction methods is not especially straightforward, however. Many different specialized image feature extraction methods have been developed, and it would be desirable to have these combined in a freely available library, as was attempted in this study. Therefore, our work on the *ImgPheno* library is on-going. For example, in an effort to normalize images prior to feature extraction, we implemented some colour image enhancement methods (developed by [25]). However, most images used were already of good quality, and performing hue-preserving linear transformation with maximum contrast often did not result in an enhanced image, and identification accuracy was not significantly improved. In other cases, where discrete patterns and shapes are more important than colour, additional algorithms might be of use. For example, in extracting features that capture venation patterns such as on leaves or insect wings, algorithms for increasing contrast and for detecting edges and intersections may yield good results.

The *ImgPheno* library was developed in Python, but to make such a library more suitable for high-throughput purposes, it might be useful to port it to a lower level programming language that can easily be interfaced to other programming languages (e.g. C++). It might even make more sense to extend existing computer vision libraries with our algorithms. OpenCV, which *ImgPheno* uses, is an open-source library that already implements a range of image feature (detection) algorithms, and such a library could easily be contributed to with new algorithms. As a bonus it already comes with interfaces to many other programming languages.

The prototype of the *OrchID* web application renders web pages that are adaptive to different screens, such that the interface looks good both on desktop computers and handheld devices. However, serving the framework in this way precludes access to advanced features in handheld devices, such as direct access to the camera or to GPS functionality (useful for logging species occurrences, or for narrowing down the possibilities during identification, as done by the Leafsnap app [26]). To enable such functionality, the web application is undergoing development to have a JSON API so that clients such as native phone apps can be developed. As such, the web application is not yet publicly accessible. However, readers interested in trying out its current functionality can do so quite easily on any UNIX-like operating system, installation consists of a single terminal command and no separate web server is required, as noted in the documentation. Public hosting of the web application on servers at Naturalis Biodiversity Center is planned for 2016.

## ACKNOWLEDGEMENTS

This project was made possible by an internal grant for application-oriented research provided by Naturalis Biodiversity Center. We are thankful to the following photographers and institutions for allowing us to use their photographs for this research: Alpine-Garden.com, Crustacare, GoreOrchids.com, Greentours, Hortus botanicus of Leiden University, Orchi, Orchids.com, Vermont Ladyslipper Company, C. Riley, D. H. Baptista, R. Barkalow (First Rays Orchids), P. C. Brouwer, M. Bürki (Botanischer Garten Bern), J. Chang, J. Claessens, M. Costea, P. J. Cribb and the Swiss Orchid Foundation, J. W. Dougherty, N. and L. Dusdieker, K. Forester, M. Günther, J. J. Harrison, R. Hella, E. Hunt, D. Judge, Rick H., R. Klinge, R. Labbe (Planteck), D. Llewelyn, F. Mah, P. Mannens, J. Marcotte, D. Panczyk, R. Parsons, H. Perner, D. Potter-Barnett, P. Quintana, M. Rosim, E. S. Saulys (Connecticut Orchid Society), P. Tremain, R. Takase, R. van Vugt, Y. Wang, D. Winkler.

## SUPPLEMENTARY MATERIALS

**Supplementary Figure 1.** Entity-relationship diagram of the metadata database used for storing taxonomic information for a collection of reference photos.

**Supplementary Table 1.** Photos available for this study.

